# Integrated proteasomal and lysosomal activity shape mTOR-regulated proteome remodeling

**DOI:** 10.1101/2024.07.20.603815

**Authors:** Danica S. Cui, Samantha M. Webster, Joseph H. Davis

## Abstract

The crucial growth regulator mTOR is suppressed during nutrient limitation, which reduces protein synthesis and activates the ubiquitin-proteasome system (UPS) and lysosomal degradation pathways. Whereas these pathways have been extensively studied individually, their integrated dynamics, including the interplay between protein synthesis and degradation, and the coordination between lysosomal and UPS pathways, remain underexplored. Here, we couple stable isotope pulse-labeling and mass spectrometry to quantify and kinetically model proteome dynamics following mTOR inhibition in cultured human cells. Using a combination of genetics and pharmacological inhibitors, we identify proteins strictly degraded by one pathway, those that undergo multimodal degradation, and others that can flexibly access the proteasome or lysosome subject to the availability of either. Our data resource, comprised of ∼5.2 million proteomic measurements, reveals that the UPS and lysosomal pathway operate with disparate kinetics, and highlights the rapid nature of lysosomal degradation. Additionally, we observe that cells coordinate the synthesis and degradation of translation initiation and elongation factors, leading to preferential synthesis from key classes of mRNA transcripts. Taken together, this work illuminates the complex, integrated pathways influencing proteostasis when mTOR is inhibited, provides a rich resource detailing the kinetics of protein synthesis and degradation, and establishes a robust methodology for measuring proteome dynamics on a per-protein basis in the context of cellular stress.

## INTRODUCTION

The mammalian target of rapamycin (mTOR) signaling pathway sits at the intersection of cellular anabolism and catabolism, where key complexes bearing the mTOR kinase (mTORC) regulate the protein synthesis and degradation pathways that govern cellular growth in response to environmental stimuli such as growth factors and nutrient levels (Thoreen, Kang *et al*. 2009, Liu and Sabatini 2020). mTOR promotes protein synthesis by activating cap-dependent translation, primarily through the phosphorylation of p70 S6 kinase 1 (S6K1) and the eukaryotic initiation factor 4E-binding proteins (4E-BP1) (Choo, Yoon *et al*. 2008). Conversely, adverse environmental conditions, including starvation, inhibit mTOR, resulting in translational suppression (Inoki, Zhu *et al*. 2003, Thoreen, Chantranupong *et al*. 2012). Notably, mTOR inhibition blocks translation of mRNAs with a 5’terminal <u>o</u>ligo<u>p</u>yrimidine (5’TOP) motif (Thoreen, Chantranupong *et al*. 2012), which is commonly found in ribosomal proteins and pro-translation factors (Kimball 2002, Iadevaia, Caldarola *et al*. 2008). More specifically, this suppression is thought to occur via dephosphorylation of 4E-BP1 following mTOR inhibition, which allows 4E-BP1 to bind to eIF4E, thereby inhibiting its interaction with eIF4G1, leading to inhibition of 5’cap-dependent translation initiation (Yanagiya, Suyama *et al*. 2012, Cockman, Anderson *et al*. 2020). Notably, Weber and colleagues have described a translation initiation pathway that is independent of eIF4E, instead relying on the eIF4G1 structural homolog DAP5 (Weber, Kleemann *et al*. 2022), which is also known as NAT1 or eIF4G2. This observation raises the possibility of differential translation of specific genes in response to mTOR inhibition.

Catabolically, mTOR inhibition causes an increase in protein degradation via activation of the ubiquitin-proteasome system (UPS), as well as lysosomal degradation via the induction of autophagy (Zhou, Tan *et al*. 2013, Zhao, Zhai *et al*. 2015). Thus, mTOR serves as a central modulator in the maintenance of cellular proteostasis, helping to adapt the proteome to varying environmental cues. In mammalian cells, proteins are often targeted for degradation upon ubiquitinylation through the activity of specific ubiquitin ligase proteins (Hershko and Ciechanover 1998). By controlling the ubiquitinylation pattern on targeted proteins, cells are thought to install a “ubiquitin code” that effectively differentiates substrates destined for lysosomal degradation from those targeted to the UPS (Komander and Rape 2012). Notably, mTOR inhibition stimulates a rapid increase in protein ubiquitination across wide swaths of the proteome (Zhao, Zhai *et al*. 2015). Such stimulated ubiquitylation can lead to increased proteolysis by the UPS (Lee and Goldberg 2022), providing one key link between nutrient sensing and proteolysis. Moreover, when active, mTOR inhibits autophagy by phosphorylating the ULK1 complex, a key component involved in autophagy initiation (Jung, Ro *et al*. 2010). When mTOR is inhibited, this repression is relieved and autophagy is stimulated. This leads to the formation of double-membraned vesicles known as autophagosomes that engulf targeted substrates and eventually fuse with lysosomes, where the substrates are hydrolyzed (Levine and Kroemer 2008). Canonical autophagy relies on the core autophagy related protein 7 (ATG7), and it represents the predominant form of autophagy activated under nutrient stress (Mizushima, Yoshimori *et al*. 2011), although non-canonical autophagy pathways that do not rely on ATG7 have been described (Nishida, Arakawa *et al*. 2009).

The interaction between the UPS and lysosomal degradation pathways is crucial for maintaining proteostasis and these pathways are known to impact one another (Kocaturk and Gozuacik 2018). However, the specifics of the interplay between them remain unresolved. For instance, pharmacological or genetic inhibition of the proteasome can lead to the upregulation of autophagy-related genes (Zhu, Dunner *et al*. 2010, Kyrychenko, Nagibin *et al*. 2014), hinting at a possible compensatory mechanism. The exact substrates targeted by such postulated compensatory regulation have yet to be identified. Additionally, there are conflicting reports regarding the effect of autophagy inhibition on proteasome activity, with some reporting reduced proteasomal activity (Korolchuk, Mansilla *et al*. 2009), and others observing no effect (Komatsu, Waguri *et al*. 2006). Notably, these studies rely primarily on indirect observations, such as changes in autophagy protein levels or ubiquitination status, and fail to conclusively determine how substrates are prioritized for degradation, particularly under stress conditions such as nutrient limitation.

Here, by coupling stable-isotope pulse labeling to quantitative mass spectrometry (Chen, Sperling *et al*. 2012, Telusma et al. *in preparation*), we independently monitor changes in the rates of protein synthesis and degradation across wide swaths of the proteome in cultured human cells. We applied this method in the context of genetic and pharmacological interventions to probe interactions between the UPS and lysosomal systems. Combined with kinetic modeling, this work provides a rich, quantitative resource detailing how the cell employs the complementary UPS and lysosomal pathways to modulate protein degradation in response to stress. We further detail how this resource can be mined to uncover preferred and alternative modes of degradation for specific proteins and how these pathways interact to fine-tune protein levels in response to nutrient starvation.

## RESULTS

To globally monitor protein dynamics in cultured human cells, we developed and applied a SILAC pulse-labeled mass spectrometry (PL-MS) workflow (**Figure 1A**). Briefly, HeLa cells were adapted to suspension culture bearing a “light” (L) isotope medium consisting of ^12^C-, and ^14^N-labeled arginine (R) and lysine (K). To initiate our measurements, we transitioned the cells to a “heavy” (H) isotope medium bearing ^13^C- and ^15^N-labeled R and K and added either the mTOR inhibitor torin1 (torin), to mimic nitrogen starvation (Thoreen, Kang *et al*. 2009), or DMSO, as a control. When coupled to whole cell proteomics, this rapid media switch allowed us to differentiate between degradation of light-labeled proteins and the synthesis of new heavy-labeled proteins. Pulse-labeling time-courses for torin and DMSO conditions were performed in triplicate, and we combined, normalized, and filtered the data from all replicates (**Supplementary Figure 1**) to generate kinetic profiles for 2,694 proteins under torin treatment, and 2,550 proteins in the control conditions (see Methods). As expected for mass spectrometry-based assays, our proteome coverage was enriched for highly abundant proteins (**Supplementary Figure 2**).

**Figure 1.**
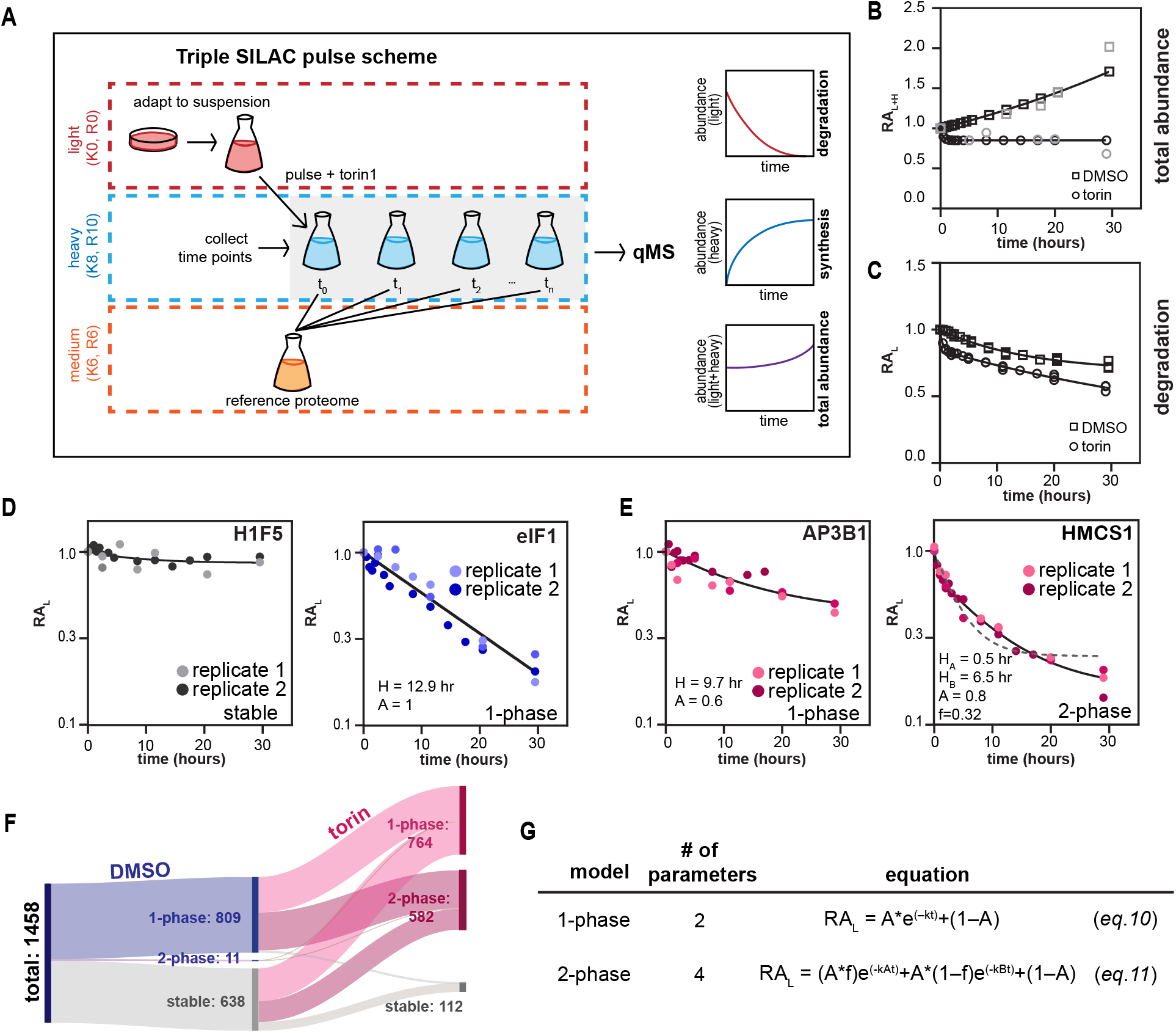
Modeling the kinetics of protein turnover through pulsed-SILAC coupled to quantitative MS. **(A)** A depiction of our method that couples pulsed-SILAC with LC-MS/MS quantitation in human cells cultured in suspension. **(B)** Global measures of total protein abundance. For each time point, the median total protein abundance relative to that at t=0 (RA_L+H_), as measured by quantitative mass spectrometry (qMS), is plotted in black for DMSO-(squares) and torin-treated cells (circles). The relative total protein mass, as measured in bulk by a Pierce A_660_ colorimetric assay, is plotted in grey. Data from qMS measurements are fit to a single exponential. **(C)** Global measure of protein degradation. For each time point, the median light abundance relative to that at t=0 (RA_L_) is plotted for DMSO and torin-treated cells. Data are fit to *eq. 10* for DMSO and *eq. 11* for torin-treated cells (see Methods). **(D)** Exemplar kinetic traces observed in the DMSO-treated cells corresponding to stable proteins (left; H1F5) and proteins that are degraded (right; eIF1), plotted on a semi-log axis. Data are fit to *eq. 10*. Varying shades denote independent replicates. **(E)** Exemplar kinetic traces observed in torin-treated cells, including proteins whose degradation kinetics are well fit (see Methods) by a 1-phase decay model (left; AP3B1; *eq. 10*), or a 2-phase decay model (right; HMCS1; *eq. 11*). For comparison, the best fit achieved by a 1-phase decay model is depicted with a dashed gray line (right). Varying shades denote independent replicates. **(F)** Sankey plot summarizing changes in the distribution of proteins fit by different kinetic models in DMSO- and torin-treated cells. Number of proteins fit by each kinetic model annotated. **(G)** Summary of equations and model parameters for 1-phase and 2-phase decay models.

In control cells, we observed a time-dependent accumulation of total protein mass as measured by <u>q</u>uantitative <u>m</u>ass <u>s</u>pectrometry (qMS) that was consistent with both a bulk colorimetric estimate of total protein abundance and observed cellular growth (**Figure 1B**). Upon torin treatment, cell growth ceased, and we observed a time-dependent decrease in total protein abundance, as previously observed (Thoreen, Kang *et al*. 2009). Analysis of heavy isotope incorporation, which quantified *de novo* protein synthesis, revealed a two-fold reduction in the median protein synthesis rate upon torin treatment (**Supplementary Figure 3**).

### mTOR inhibition modulates translational control in a gene-specific manner

At the individual mRNA level, we observed that torin treatment led to transcript-specific translational regulation. Specifically, the translation of mRNAs bearing the 5’-TOP motif was suppressed, with up to 10-fold repression of some genes observed (**Supplementary Figure 4A**). In contrast, the translation of other proteins was unaffected by torin treatment (**Supplementary Figure 4B**). To assess the impact of transcription on these observations, we re-analyzed RNA-seq data from a prior study on HEK293T cells treated with torin for 24 hours (Park, Reyna-Neyra *et al*. 2017), and we found no notable alterations in mRNA levels for these proteins (**Supplementary Figure 4C**). This was consistent with transcript-specific regulation of translation rather than alterations in mRNA abundance.

### mTOR inhibition dually activates lysosomal and proteasomal pathways

Having established the impact of torin treatment on protein synthesis, we next explored the role of degradation in remodeling of the proteome. Upon inspection of the light isotope channel, which reported on protein degradation kinetics, we observed a global increase in the extent of protein degradation in the torin-treated condition compared to the control (**Figure 1C**). At the individual protein level, we observed diverse kinetic behaviors. Some protein degradation profiles were well fit by a 1-phase (*i*.*e*. single exponential) decay model, suggesting a uniform degradation process, whereas others were better fit with a 2-phase model, consistent with the presence of multiple concurrent degradation processes with distinct rates of degradation, or multiple pools of a given protein with differential stability. For example, we hypothesized that lysosomal and proteasomal degradation might target substrates in parallel but at different rates, or that a given protein could bear variable post-translational modifications or localizations that impacted stability. To model these diverse profiles, we employed an integrated data analysis workflow (**Supplementary Figure 1**, see Methods) resulting in kinetic models for degradation of 1,658 proteins in the DMSO-treated condition and 2,015 proteins in the torin-treated condition.

At the individual protein level, our analysis revealed profound changes in the distribution of 1-phase and 2-phase degradation profiles upon torin treatment. Indeed, in the control condition, 55% of the degradation kinetic profiles were well fit by the 1-phase model, whereas 44% of these profiles were effectively stable over the 29-hour time-course (**Figure 1D**). In contrast, upon torin treatment, most monitored proteins were degraded, with observed kinetic profiles consistent with both 1-phase (51%) and 2-phase (41%) models of degradation (**Figure 1D-E**). We also identified a small fraction (8%) of measured proteins that escaped degradation, which indicated that the reduction in light-labeled proteins we observed was not a result of cell lysis or other global losses of protein occurring during drug treatment sample preparation.

Previous studies have documented the activation of both lysosomal and proteasomal pathways upon torin treatment (Zhou, Tan *et al*. 2013, Zhao, Zhai *et al*. 2015). As such, we hypothesized that some of the biphasic kinetics observed under torin treatment could result from the combined contributions of proteasomal and lysosomal degradation pathways. To investigate this hypothesis, and to determine the respective roles of these pathways in protein degradation, we collected additional time-courses in the presence of either torin and bortezomib (t/bort), which inhibits the proteasome (Chen, Frezza *et al*. 2011); or torin and bafilomycin A (t/baf), which inhibits lysosomal acidification and activity (Mauvezin and Neufeld 2015). We additionally measured protein degradation following co-treatment with torin, bafilomycin A, and bortezomib (t/bort/baf). We focused our analyses on the 577 proteins that were degraded under torin treatment, but were stable in t/bort/baf conditions, as these substrates were likely to be either lysosomal or UPS targets (**Figure 2A**). Kinetic modeling of the observed degradation profiles revealed three groups with distinct behaviors (**Figure 2B**). Group A proteins (n = 205) exhibited 2-phase degradation under torin treatment but were fit by a 1-phase model when treated with either t/baf or t/bort. Group B proteins (n = 11) ere well fit by the 1-phase model under torin and t/ bort treatment, but were stable when treated with t/baf. Group C proteins (n = 145) were well fit by a 1-phase model under all three treatment conditions, but were stable in the t/bort/baf treatment. The remaining 108 proteins exhibited relatively small numbers of the remaining condition-dependent combinations of stable, 1-phase, and 2-phase behavior. As discussed in detail below, we hypothesized that groups A-C delineated three key categories of proteome dynamics of interest: proteins undergoing a combination of lysosomal and proteasomal degradation in torin-treated cells (group A); proteins exclusively degraded by lysosomes (group B), and proteins that flexibly access both lysosomal and proteasomal degradation pathways (group C).

**Figure 2.**
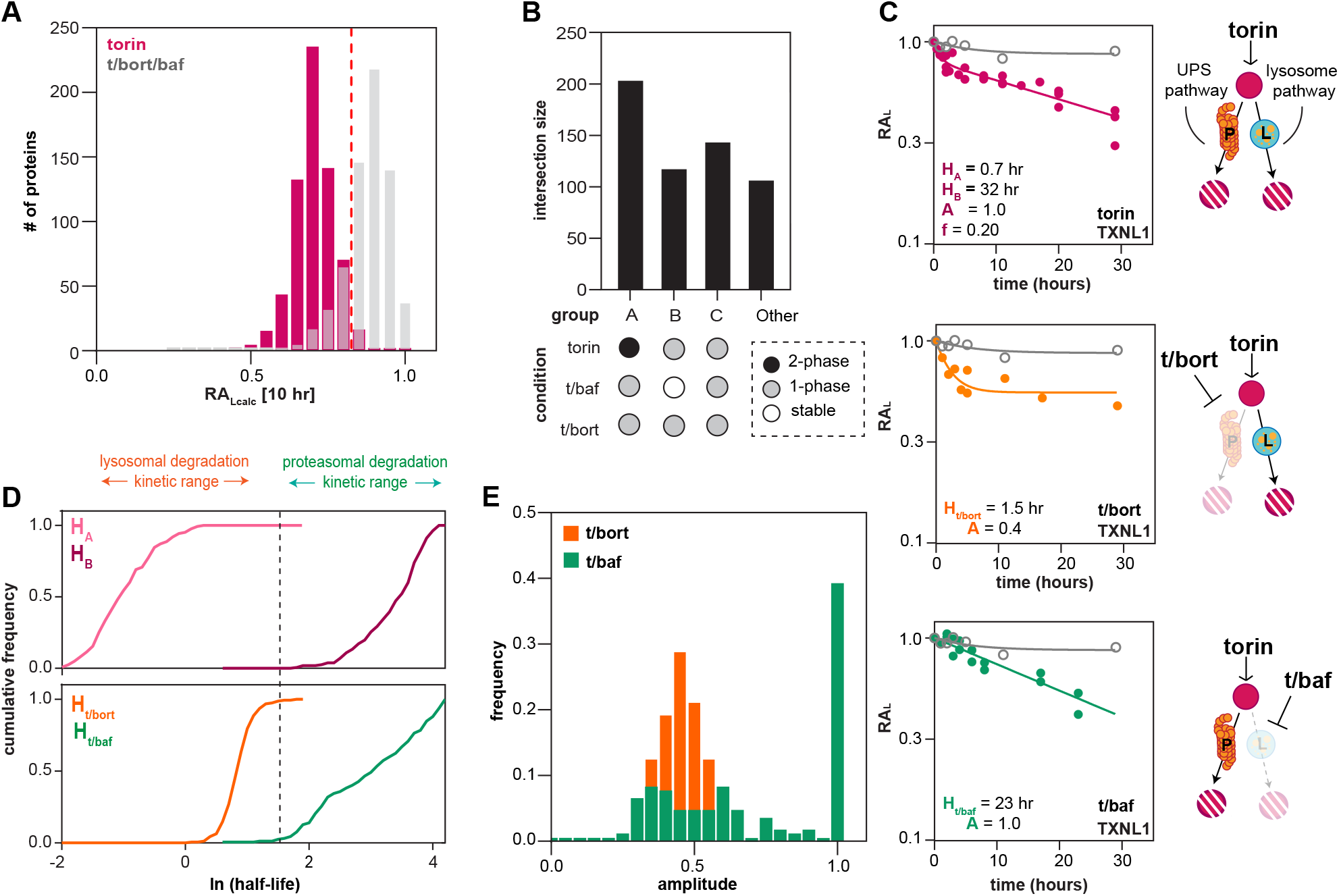
Lysosomal and proteasomal degradation pathways exhibit disparate kinetic profiles. **(A)** Histogram of protein counts vs. the relative abundance as calculated from fit kinetic profile(RA_Lcalc_) at 10 hours post-treatment with torin (pink) or torin/bort/baf (grey). A cutoff indicated by the dashed line (>0.85 RA_Lcalc_) delineates proteins considered stable under torin/bort/baf. These proteins were used for subsequent analysis. **(B)** An UpSet plot categorizing proteins based on their kinetic characteristics in response to treatment with torin, t/baf, and t/bort. **(C)** Kinetic degradation profiles for TXLN1 in the torin (top), t/bort (middle), and t/baf (bottom) conditions, with the t/bort/baf treatment overlaid in grey for comparison. **(D)** A plot of the cumulative frequency of half-lives for group A proteins treated with torin (top), and those treated with t/baf and t/bort (bottom). **(E)** Distribution of degradation amplitudes across all group A proteins under either t/baf and t/bort treatment regimes.

### Lysosomal and proteasomal degradation profiles are kinetically distinguishable

To interpret the biphasic labeling kinetics observed for group A proteins, we first inspected the fit kinetic parameters for exemplar proteins such as thioredoxin like protein 1 (TXNL1). There, we noted that the 2-phase parameters observed upon torin treatment could be decomposed into the isolated 1-phase parameters measured upon lysosomal or proteasomal inhibition (**Figure 2C**). Specifically, treatment with torin produced a rapid burst phase (pathway 1) for degradation with a half-life of 0.7 hours (H_A_), and a slower but persistent second phase (pathway 2) with a half-life of 32 hours (H_B_). Under t/baf or t/bort treatment conditions, the kinetics resolved to a single phase with observed half-lives similar to each of the two states observed in torin treatment. We noted that the kinetic characteristics of each pathway were distinct, with the lysosomal pathway executing more rapid TXLN1 degradation and the proteasome degrading a larger fraction of the total TXLN1 substrate, albeit at a reduced rate.

Given the expansive nature of this dataset, we next asked whether the distinct kinetic features observed for TXLN1 were common across the proteome. Briefly, we inspected the cumulative distribution of the fast phase and slow phase half-lives for Group A proteins under torin treatment, and compared those to the single-phase half-lives observed upon t/bort or t/baf treatment. Consistent with our analysis of TXLN1, we observed distinct timescales on which the fast (H_A_) and slow (H_B_) processes operated, with more rapid single-phase degradation upon proteasomal inhibition than upon lysosomal degradation. Specifically, we observed half-lives under t/bort treatment that were approximately five times shorter than those under t/baf treatment, consistent with rapid lysosomal degradation (**Figure 2D**). We also inspected the amplitude of each fit under t/bort and t/baf treatments, which reported on the fraction of each substrate that was degraded, and found that proteasomal inhibition limited the extent of substrate degradation more than lysosomal inhibition (**Figure 2E**). Taken together, these results suggested that the proteasome, although slower, was a more processive degradative pathway than the lysosome, whose degradative capacity was rapidly exhausted.

### ATG7 is required for some, but not all, lysosome-dependent protein degradation

To understand the contribution of canonical autophagy versus other lysosomal-targeting pathways to degradation, we used a previously reported ATG7 knock-out (ATG7Δ) HeLa cell line (Abudu, Pankiv *et al*. 2019), which ablates canonical autophagy by blocking conjugation of LC3/ GABARAP family proteins to the autophagosomal membrane (Mizushima, Sugita *et al*. 1998). Kinetic analysis of torin-treated cells showed a shift in the kinetic profiles of protein degradation: 87% of the proteins that displayed 2-phase kinetics in torin-treated wild-type cells transitioned to single-phase kinetics in ATG7Δ cells (**Figure 3A**), emphasizing the role of ATG7-dependent pathways in mediating torin-induced biphasic degradation.

**Figure 3.**
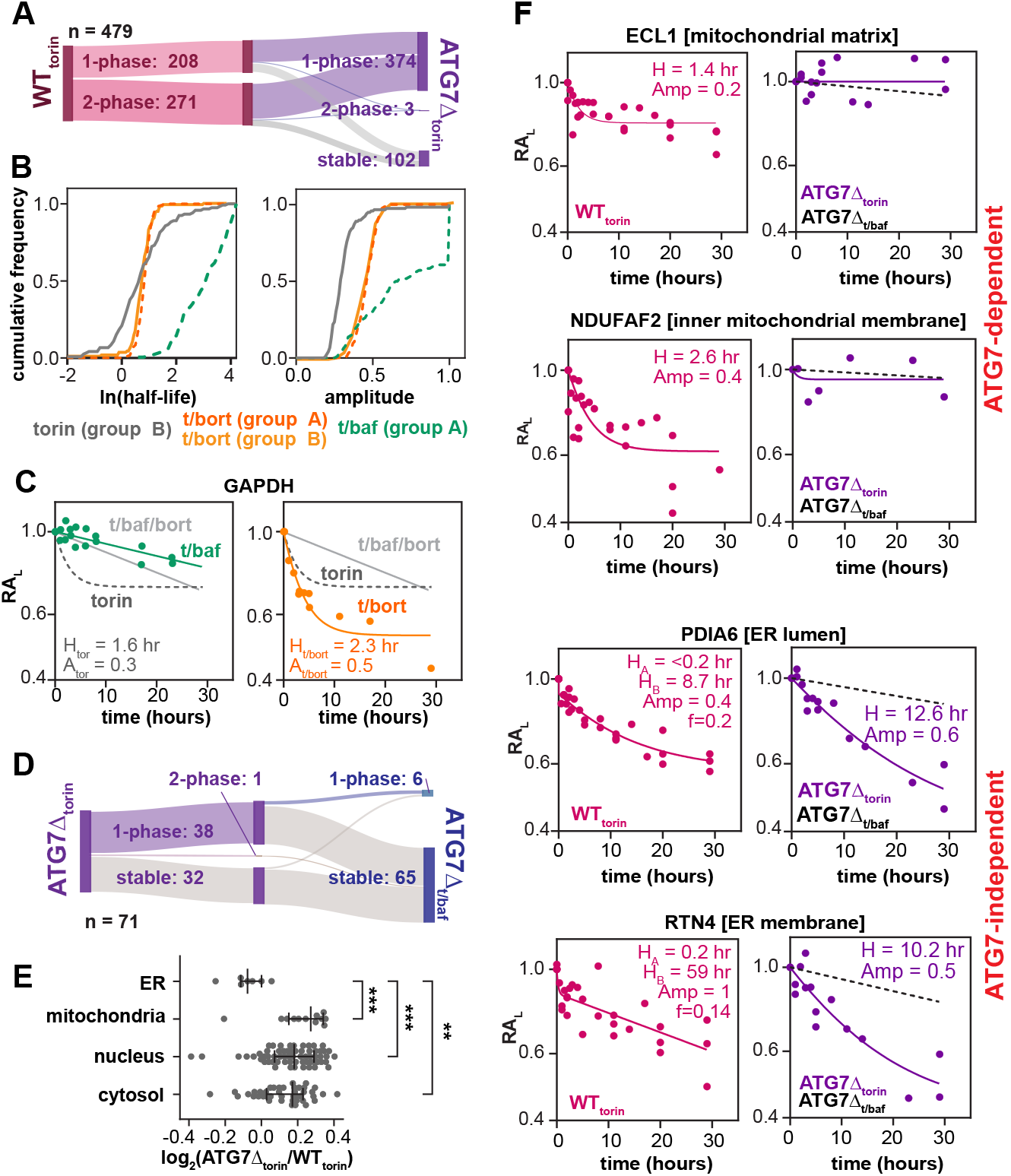
ATG7 is required for some, but not all, lysosomal proteolysis. **(A)** Sankey plot depicting kinetic classification of wild-type and ATG7Δ cells treated with torin. **(B)** Cumulative frequency plots comparing fit 1-phase kinetic degradation parameters of group A (n=205) and group B (n=119) proteins upon torin, t/baf and t/bort treatment. Note that t/baf treatment ablated degradation of group B proteins. **(C)** Exemplar kinetic curves of GAPDH, a known lysosomal substrate, shown on a semi-log axis. Data fit to 1-phase degradation model (*eq. 10*), with fit parameters shown. **(D)** Sankey plot of group B proteins (n=71) reflecting the kinetic classification of ATG7Δ cells treated with torin or t/baf. **(E)** A scatter plot depicting the fold-change in protein abundance in torin-treated wild-type vs. ATG7Δ cells. For each protein, abundance was determined using fit 1-phase kinetic model, evaluated 10-hours post-torin treatment, and proteins (individual points) are grouped by sub-cellular localization as annotated in Thul *et al*. 2017. Analysis was focused only on proteins that were stable when treated with t/bort/baf. The median and interquartile range are depicted in black. One-way ANOVA analysis with statistical significance (** p< 0.005, *** p<0.0005) noted. **(F)** Kinetic traces of proteins whose degradation was ATG7-dependent (top) or ATG7-independent (bottom). Kinetic parameters fit using the 1-phase (top) and 2-phase (bottom) degradation models noted.

Next, we inspected the torin-dependent kinetic profiles of group B proteins, which were degradation-resistant when treated with t/baf, suggesting lysosomal-targeting of these proteins. Consistent with this model, degradation profiles for group B proteins closely resembled the lysosome-dependent degradation phase assigned to group A protein degradation kinetics (**Figure 3B**). This conclusion was also supported by inspection of the kinetic trace of a known lysosomal substrate, glyceraldehyde-3-phosphate dehydrogenase (GAPDH) (Aniento, Roche *et al*. 1993), whose torin-dependent degradation was ablated in the t/baf but not in the t/bort treatment regime (**Figure 3C**). Surprisingly, among group B proteins, we found that only 49% relied on ATG7 for lysosome-dependent degradation (**Figure 3D**), underscoring the existence of ATG7-independent lysosomal degradation pathways (Mejlvang, *et al*. 2018).

As ATG7-dependent autophagy has been previously linked to degradation of organelles (Komatsu, Waguri *et al*. 2005), we next examined the subcellular localization of substrates and compared their degradation kinetics in our WT_torin_, ATG7Δ_torin_, and ATG7Δ_t/baf_ datasets. This analysis revealed a strong ATG7-dependence on the degradation of mitochondrial proteins, especially those in the mitochondrial matrix (*e*.*g*. ECI1), and in the inner membrane (*e*.*g*. NDUF2). Conversely, proteins located in the ER were largely degraded in an ATG7-independent manner (**Figure 3E-F**). Overall, these findings suggested that ATG7-dependent autophagy plays an important role in lysosomal protein degradation, but highlighted that this effect is not uniform as ATG7 was dispensable for lysosomal degradation of many ER-localized proteins.

### UPS and lysosomal degradation pathways are mutually compensatory

We next investigated the dynamics of group C proteins, which were well fit by the 1-phase degradation model in torin, t/baf, and t/bort treatment regimes, but were stable in cells treated with t/bort/baf. Inspection of the fit parameters in torin-treated cells revealed distinct protein sets – one consistent with relatively rapid but incomplete lysosomal degradation, and a second consistent with slower, but more persistent proteasomal degradation. For example, parameters used to fit degradation of Heterogeneous nuclear ribonucleoprotein F (HNRNPF), a known proteasomal substrate (Chabes and Thelander 2000, Baugh, Viktorova *et al*. 2009, Koumbadinga, Mahmood *et al*. 2015), in torin-treated cells were most consistent with proteasomal degradation. In line with this model, addition of bafilomycin A minimally impacted HNRNPF’s degradation kinetics. Intriguingly, when proteasomal activity was inhibited, HNRNPF was still degraded, though with distinct kinetics that were consistent with lysosomal degradation (**Figure 4A**).

**Figure 4.**
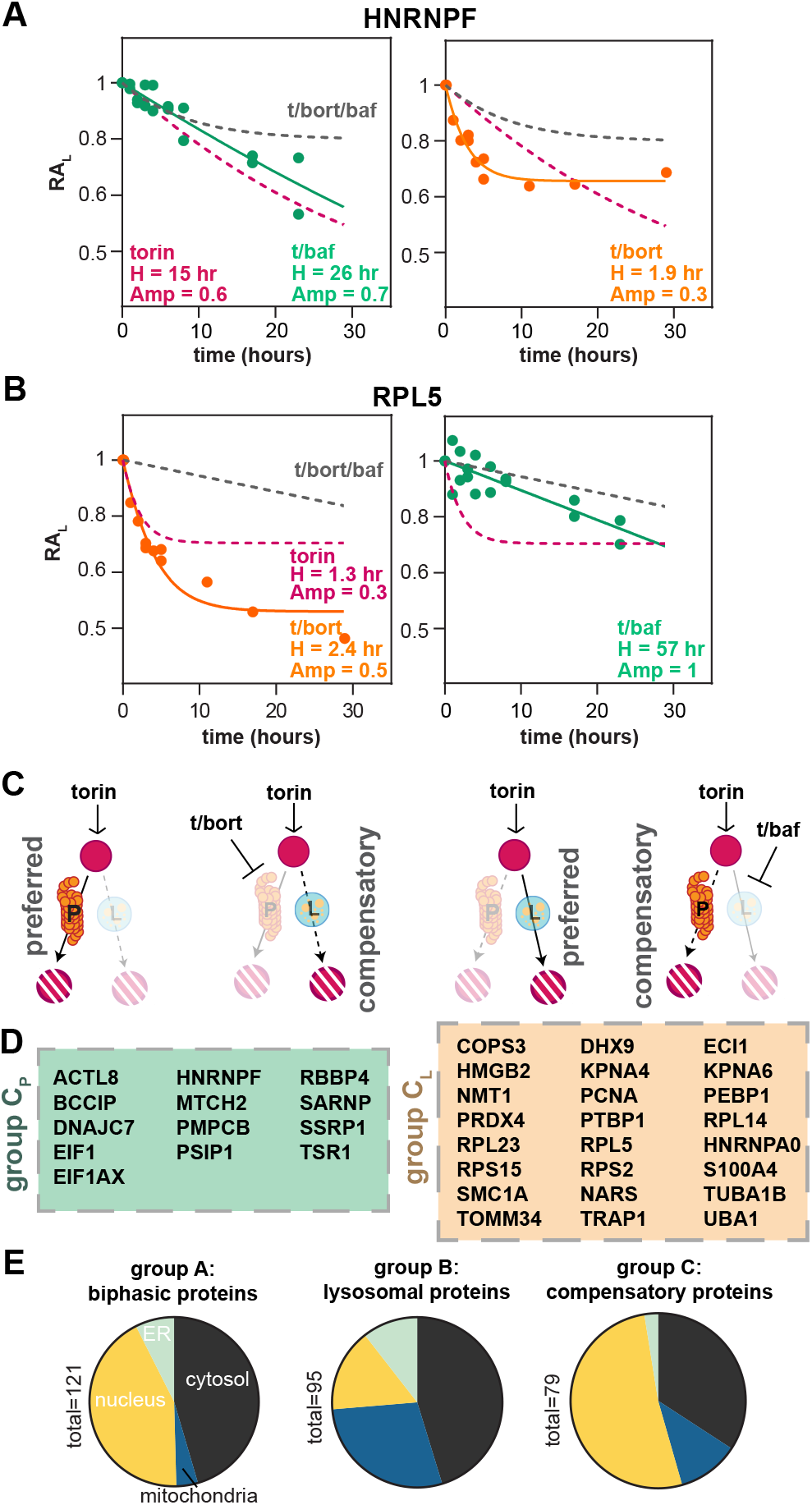
Kinetic analyses resolve compensation between proteasomal and lysosomal degradation pathways. **(A)** Degradation profiles of HNRNPF, an exemplar proteasomal substrate that exhibits lysosomal compensation in t/bort treatment conditions. Data are fit to the 1-phase model (*eq. 10*), with fit parameters for each treatment regime noted. **(B)** Degradation profiles of RPL5, an exemplar lysosomal substrate that exhibits proteasomal compensation in t/baf treatment conditions. Data are fit as in (A). **(C)** Cartoon model of preferred and compensatory degradation pathways for group C_P_ (left) and group C_L_ (right) proteins. **(D)** Catalog of group C_P_ (left) and group C_L_ (right) proteins that are preferentially degraded by a single pathway in torin-treated cells, but utilize the compensatory pathway upon pharmacological (t/bort, or t/baf) inhibition of the preferred pathway. Kinetic fits for each protein are provided in Supplementary Figure 5. **(E)** Depiction of subcellular localization of group A, B, and C proteins, as annotated in Thul *et al*. 2017.

Analogously, torin-dependent degradation kinetics for ribosomal protein L5 (RPL5) and multiple other ribosomal proteins were most consistent with lysosomal degradation, and addition of bortezomib did not inhibit this degradation (**Figure 4B**). Notably, lysosomal degradation of ribosomes has been previously reported (Wyant, Abu-Remaileh *et al*. 2018), though An *et al*. showed that such degradation was independent of canonical selective autophagy (An and Harper 2018, An, Ordureau *et al*. 2020). Interestingly, RPL5 was still degraded in the t/baf treatment regime, and did so with kinetics most similar to proteasomal substrates.

Taken together, these analyses of HNRNPF and RPL5 suggested the existence of: 1) a preferred degradative pathway for each substrate, which could be lysosomal or proteasomal; and 2) the existence of compensatory pathways that supported degradation upon inhibition of the preferred pathway. We next inspected the fit amplitude and rate parameters for all group C proteins in cells treated with torin, t/baf, and t/bort, and annotated each protein’s preferred and alternative degradative pathway based on these condition-dependent fit parameters (**Supplementary Figure 5**). We labeled proteins preferentially degraded by the proteasome as group C_P_, and those preferentially degraded by the lysosome as group C_L_, noting that the lysosomal and proteasomal pathways could effectively “compensate” for one another upon inhibition of the preferred pathway (**Figure 4C-D**). As predicted by this compensatory model, proteins within group C_P_ were robustly degraded in ATG7Δ cells treated with torin or t/ baf, and proteins from group C_L_ exhibited kinetic profiles consistent with proteasomal degradation in ATG7Δ cells (**Supplementary Figure 6**). Finally, we inspected the subcellular localization of the group A-C proteins, finding that groups using both UPS and lysosomal degradation pathways (*i*.*e*. groups A and C) were enriched in proteins localized to the nucleus, hinting at an intricate interplay of UPS and lysosomal degradation pathways within this compartment (**Figure 4E**).

### mTOR inhibition disproportionally impacts levels of key translation factors

Given mTOR’s well-documented regulation of protein synthesis and the decrease in translation we observed upon torin treatment, we next asked how torin impacted the degradation of ribosomal proteins and translation factors. We observed that the factors were, on average, degraded to a larger extent than the core ribosomal protein components (**Figure 5A**), which we speculated could facilitate a more rapid translational response than could be achieved through widescale ribosomal degradation. Extending this hypothesis, we next asked whether the total abundance of specific translation factors was differentially regulated in response to torin. We observed a 40% decrease in the levels of eukaryotic elongation factor 2 (eEF2), which we expected would slow translation and decrease polysome formation (Schieweck, Ciccopiedi *et al*. 2023). In contrast, the level of translation initiation factor 6 (eIF6), an inhibitor of small and large subunit association (Jaako, Faille *et al*. 2022), increased by 20% (**Figure 5B-C**). This shift in the relative proportions of a pro-elongation factor (eEF2), and an anti-association factor (eIF6) hinted at an mTOR-dependent mechanism to limit the assembly and translational capacity of mature 80S ribosomes. This hypothesis was consistent with ribosome sedimentation profiles, which displayed a large 60S subunit peak under torin compared to DMSO treatment, as well as an accumulation of 80S monosomes and a reduction in polysomes (**Figure 5D**). We also explored if changes in the abundance of initiation factors could explain the mRNA-specific translation we had observed. We specifically focused on the alternative 5’cap-dependent translation initiation factors DAP5 and EIF3D, as we had observed robust synthesis of EGFR (**Supplementary Figure 4B**), which depends on DAP5 for translation initiation (de la Parra, Ernlund *et al*. 2018). After 29 hours of torin treatment, we observed a reduction in the levels of canonical cap-dependent translation factors eIF4G1 and eIF4E relative to that of DAP5 and EIF3D (**Figure 5E-F**), which we expected would favor DAP5-mediated synthesis at the expense of eIF4E-dependent translation. Indeed, we observed a ∼5-fold reduction in the synthesis of mRNAs bearing 5’TOP motifs, which require eIF4E for translation. In contrast, annotated DAP5-dependent genes exhibited moderate repression (**Figure 5G**). Taken together, these analyses of proteome dynamics highlight how cells link nutrient sensing, regulated proteolysis, and translational reprogramming to remodel their proteome in response to environmental stimuli, and it provides a resource rich with testable hypotheses for future mechanistic studies.

**Figure 5.**
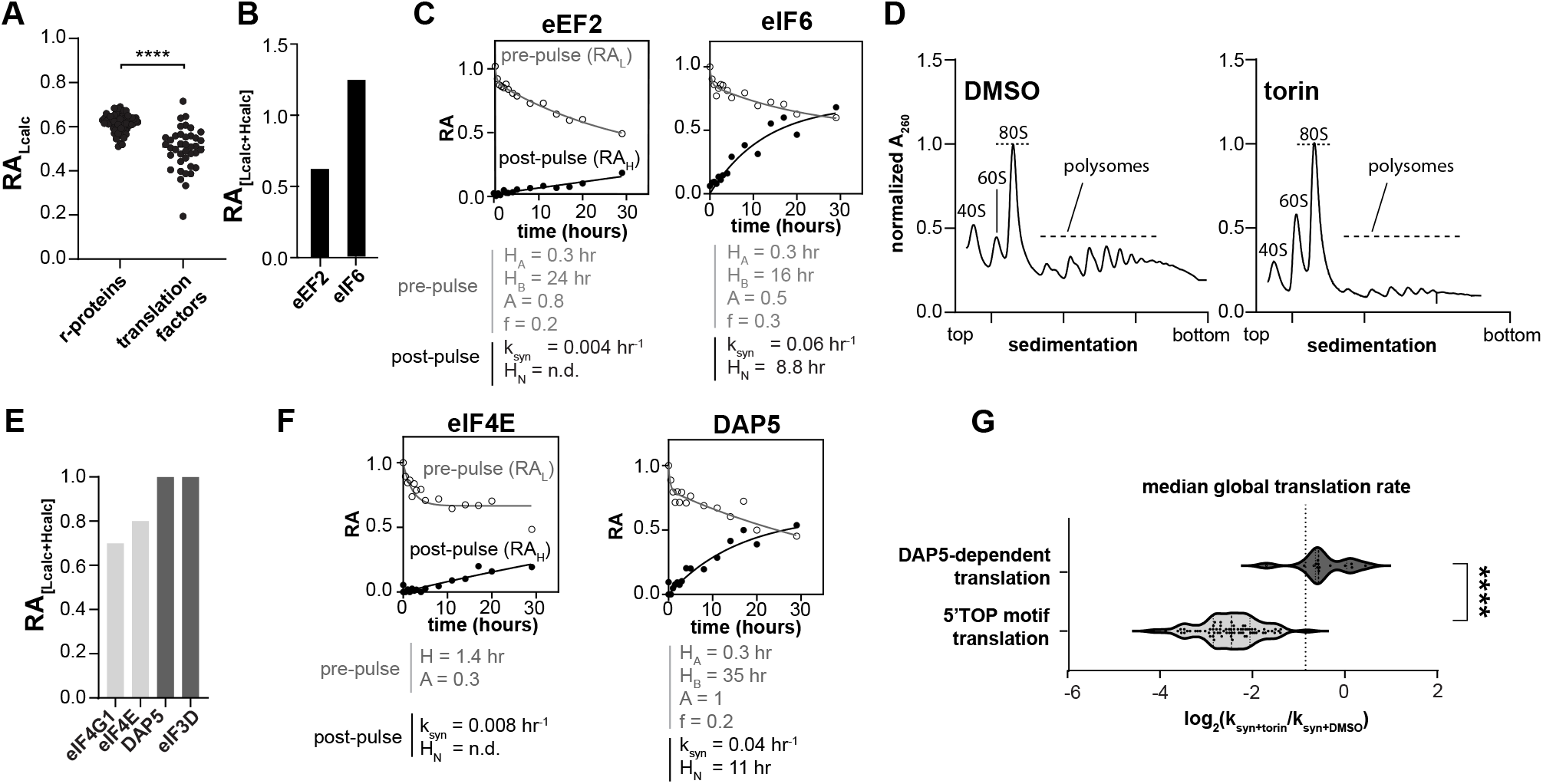
Modulation of translation factor levels drives torin-dependent translational reprogramming. **(A)** A comparison between the relative abundance (RA_Lcalc_) of r-proteins and translation factors (as annotated by gene ontology term GO:0008135) after 29 hours of torin treatment. Abundance calculated from fit of entire kinetic profile, with dots representing individual proteins. Statistical significance of the abundance change between groups was measured by a two-tailed t-test, and is noted with asterisks (p<0.0001). **(B)** Comparison of the change in the relative total protein abundance calculated from fitting of the pre-pulse (L) and post-pulse (H) kinetic profile of pro-translation (eEF2) and anti-translation (eIF6) factors. **(C)** Kinetic profiles of translation factors analyzed in (B). Protein levels are separated by isotope channel, with pre-pulse isotope abundance (open circles) and post-pulse isotope abundance (closed circles) normalized to that at t=0. Fit parameters for each kinetic profile noted following *eq. 11* (pre-pulse) and *eq. 9* (post-pulse). **(D)** Sucrose density gradient profiling of ribosomal subunits after 18 hours of treatment with 100 nM torin or DMSO treatment. For each trace, absorbance at 260 nm was normalized to that of the 80S peak. Subunits, monosomes, and polysomes noted. **(E)** Comparison of the relative total abundance of cap dependent translation factors eIF4G1 and eIF4E, and noncanonical translation factors DAP5 and eIF3D after 29 hours of torin treatment. Abundance was calculated as described in (B). **(F)** Kinetic profiles of eIF4E and DAP5 protein levels annotated as described in (C). **(G)** Distribution of torin-dependent changes in synthesis rates, as measured by fitting kinetic traces of heavy isotope label incorporation, for DAP5-dependent (dark grey) and eIF4E-dependent (light grey) transcripts. Transcript annotated for DAP5 dependent translation were retrieved from Weber et al., 2022. Transcripts with 5’TOP motifs are annotated according to Meyuhas et al., 2016. Median synthesis rate across all measured proteins noted, with statistical significance of change between groups calculated and annotated as in (A).

## DISCUSSION

mTOR plays a pivotal role in maintaining proteostasis where it modulates both protein synthesis and degradation in response to environmental signals (Thoreen, Kang *et al*. 2009, Liu and Sabatini 2020). Upon starvation, the mTOR kinase activity is suppressed, leading to a general decrease in protein translation and an acute repression of transcripts bearing 5’TOP motifs (Inoki, Zhu *et al*. 2003, Thoreen, Chantranupong *et al*. 2012, Advani and Ivanov 2019). Concurrently, mTOR suppression activates autophagic and proteasomal degradation pathways, which help cells adapt to nutrient stress by remodeling the proteome (Zhou, Tan *et al*. 2013, Zhao, Zhai *et al*. 2015). Our work expands on this understanding by quantifying the impacts of mTOR inhibition on protein synthesis and degradation on a per-protein basis across large swaths of the proteome, and by providing a vast, quantitative dataset to model the synthesis and degradation of ∼2,600 proteins across 7 strains or conditions, with ∼5.2 million total proteomic measurements reported. Our proteome-scale analysis recapitulates multiple focused studies (Zhao, Zhai *et al*. 2015, Bartolome, Garcia-Aguilar *et al*. 2017) linking mTOR to degradation of specific proteins and organelles and, further, identifies new mTOR-dependent autophagic and proteasomal substrates ripe for further study. Finally, through detailed kinetic modeling, this work has uncovered extensive collaboration and compensation between the autophagic and proteasomal degradation pathways, and it provides the requisite tools to decipher the mechanisms of this interplay.

### mTOR inhibition reprograms translation

Our work challenges the view that mTOR inhibition simply represses translation of mRNA’s bearing 5’TOP motifs (Thoreen, Chantranupong *et al*. 2012, Nandagopal and Roux 2015), and instead reveals that active translation is both downregulated and reprogrammed. First, we observed a two-fold repression in global protein synthesis as measured by reduced incorporation of mass-distinguishable pulsed amino acids upon torin treatment (**Supplementary Figure 3**), consistent with prior bulk measurements (Thoreen, Kang *et al*. 2009). Critically however, at the individual protein level, we found highly variable levels of translational suppression. Specifically, synthesis rates of transcripts containing 5’TOP motifs – a class of mRNAs known to be translationally regulated by mTOR (Thoreen, Chantranupong *et al*. 2012) – were reduced ∼4-10 fold, whereas synthesis of other transcripts was unaffected or upregulated upon mTOR inhibition (**Supplementary Figure 4B**). For example, we observed robust synthesis of EGFR, whose translation is dependent on non-canonical DAP5-dependent 5’ cap interactions (de la Parra, Ernlund *et al*. 2018). This specific observation was consistent with a more global analysis by us (**Figure 5G**) and others (Volta, Perez-Baos *et al*. 2021) showing that mTOR inhibition guides translation away from canonical eIF4E-dependent pathways and towards alternative DAP5-dependent mechanisms. As a result, translational capacity is generally directed away from synthesis of pro-growth factors such as ribosomes, and toward sensing and signaling molecules likely needed to effectively respond to the changing environment (Weber, Kleemann *et al*. 2022). Intriguingly, we newly found that mTOR inhibition alters the abundance of eIF4E and DAP5 (**Figure 5B**), providing a plausible mechanistic explanation for the observed shift in transcript selection.

We further observed a widespread torin-dependent decrease in the levels of regulatory translation factors relative to core ribosomal components. This observation implies that the changes we observed in global translation were driven in part by tuning levels of regulatory factors. Indeed, we observed differential modulation of initiation factors (*e*.*g*., eIF5B and eIF6) and the down-regulation of elongation factor eEF2, which manifested as impediments to 80S subunit joining and translation elongation as measured by sucrose gradient sedimentation assays (**Figure 5 E-G**). The relatively minor induction of ribosomal protein degradation upon starvation was similar to that observed by An *et al*. (An, Ordureau *et al*. 2020). Taken together, our findings highlight that in addition to the known mechanisms of translational regulation by mTOR, such as the phosphoregulation of S6 kinase and 4E-BPs, mTOR also indirectly modulates the abundance of translation factors to influence translational capacity and transcript prioritization, and that mTOR inhibition has a limited role in regulating the total abundance of ribosomal proteins.

### Biphasic protein degradation is triggered by mTOR inhibition

Kinetic profiling of light isotope-labeled proteins under mTOR inhibition revealed that substrates can be degraded through multiple pathways in parallel. Under torin treatment, ∼41% of the measured proteome exhibited kinetic profiles best fit by a 2-phase model (**Figure 1F**). To dissect the pathways involved, we focused our analysis on 577 proteins that were stabilized upon t/baf/bort co-treatment. Based on their distinct kinetic characteristics, we classified substrates into three groups (**Figure 2B**): Group A displayed biphasic degradation, Group B underwent lysosome-specific degradation, and Group C was comprised of proteins with compensatory degradation.

Dissecting the biphasic kinetics of group A proteins revealed a coordinated strategy that employs both the lysosome and proteasome degradation pathways in a temporally distinct manner. Specifically, we observed that the lysosomal pathway drove the rapid phase of degradation whereas the proteasome was responsible for the slower degradative phase. Such bifurcated degradation was most clearly demonstrated in the degradation kinetics of TXNL1 (**Figure 2C**), where its half-life under t/bort (H_t/bort_) treatment matched the rapid kinetic phase (H_A_) under torin treatment. Likewise, under t/baf treatment, the half-life (H_t/baf_) correlated with the slower phase (H_B_) observed in torin-treated cells. Interestingly, on a global scale, our results indicate that this biphasic degradation is dependent on ATG7 (**Figure 3B**), suggesting that the fast lysosomal degradation phase is driven by an ATG7-dependent process. This finding points to the possibility that autophagy pathways are activated as a mechanism for temporal control, thereby providing cells with the ability to respond to environmental stresses in a timely manner.

Analysis of the fit amplitudes, which represent the fraction of substrate degraded by each pathway, revealed another critical distinction between the lysosomal and proteasomal degradation routes. For the lysosomal t/baf treatment, the distribution of amplitude values centered around ∼0.5, whereas the majority of substrates under the t/bort treatment exhibited amplitudes closer to 1. This difference in amplitude suggests that the proteasome has greater substrate accessibility than the lysosome. Our findings demonstrate that cells employ a coordinated strategy involving both lysosomal and proteasomal pathways to remodel the proteome under nutrient stress, with the lysosome rapidly degrading proteins for an immediate response while the proteasome plays a more gradual role in systematically reshaping the proteome composition over a longer timeframe.

### Proteasomal and lysosomal degradation are mutually compensatory

We found our discovery of a class of proteins that can access compensatory degradation pathways (Group C proteins) notable. These proteins displayed a single-phase degradation profile under torin treatment, consistent with a preferred degradative pathway and they were stabilized by the co-inhibition of lysosomal and proteasomal pathways, implicating these pathways in their turnover. Yet, when we blocked the preferred degradative pathway, we still observed degradation and, interestingly, such degradation proceeded with a kinetic profile most consistent with the non-preferred pathway. In sum we interpret these data as evidence that these two pathways can compensate for one another, and hypothesize that such redundancy affords the cell degradative flexibility should either pathway be blocked, overloaded, or otherwise transiently unavailable.

The ability to characterize such compensatory proteins has traditionally been challenging, as measuring a single time point or assaying steady-state protein levels cannot capture the complex degradation dynamics at play. In contrast, the extensive survey of degradation kinetics provided here uncovered two distinct substrate classes: C_P_ proteins, which predominantly undergo proteasomal degradation under torin treatment, but can be processed by the lysosomal pathway upon proteasomal inhibition, and C_L_ proteins, which utilize lysosomal degradation, but can be degraded by the proteasome when lysosome acidification is blocked (**Figure 4C-D**). We hypothesize that the observed compensation could operate through common ubiquitin signaling, which plays roles in both lysosomal (Kirkin, McEwan *et al*. 2009, Stolz, Ernst *et al*. 2014) and proteasomal targeting (Collins and Goldberg 2017), with a “ubiquitin code” based on the Ub-linkage pattern thought to direct ubiquitinylated substrates appropriately (Komander and Rape 2012, Yau and Rape 2016, Lu, den Brave *et al*. 2017, Finley and Prado 2020). This raises the question of how to reconcile such a ubiquitin code with our observed compensatory degradation. One possibility is that the ubiquitin code can be bypassed upon prolonged inaccessibility of the coded pathway, allowing the alternative pathway to recognize substrates more promiscuously. Alternatively, the initial code could be removed through the activity of deubiquitinating enzymes followed by the installation of an alternatively coded mark directing the substrate to the compensatory pathway. Key questions for future study include: which of these or other possible mechanisms enable such compensatory degradation; whether recoding is actively regulated or simply the product of kinetic partitioning (Hardy and Randall 1991); and what the phenotypic consequences of a failure to compensate are. Our described approach and accompanying data resource provide key reagents in pursuing such questions.

## MATERIALS AND METHODS

### Cell culture reagents

Wild-type and ATG7Δ HeLa cells (gifts of Profs. Malene Hansen and Terje Johansen, respectively) were grown in adherent cell culture in ACC media consisting of: DMEM (GenClone #25-501) supplemented with 10% heat inactivated fetal bovine serum (Corning # 35-016-CV), 2mM Glutamax™ (Gibco # 35050061) and 1mM sodium pyruvate (Gibco #11360070). Cells were grown in suspension culture in SCC media consisting of: SILAC-Freestyle medium (a gift from the R&D Cell Biology Department, Thermo Fisher Scientific) supplemented with 1% dialyzed fetal bovine serum (Gibco #26400044), 200mg/L of L-proline (Sigma Aldrich #81709), 1.2mM of L-arginine (Sigma #11039) and 1.4mM of L-lysine (Sigma #62929). Isotopically labeled SCC media included: “medium labeled” medium with ^13^C_6_ arginine (Cambridge Isotope Labs # CLM-2247-H-0.1) and ^13^C_6_ lysine (Cambridge Isotope Labs # CLM-2247-H-0.1); or “heavy labeled” medium with ^13^C_6_, ^15^N_4_ arginine (Cambridge Isotope Labs # CLNM-539-H-0.25) and ^13^C_6_, ^15^N_2_ lysine (Cambridge Isotope Labs # CNLM-291-H-0.1).

### Pharmacological inhibitors

Torin1 (MedChem Express #HY-13003) was prepared as a 0.5mM solution in dimethyl sulfoxide (DMSO) (Corning #25-950-CQC), and used at a concentration of 100nM in cell culture. Bafilomycin A1 (Adipogen Life Sciences #BVT-0252-M001) was prepared as a 1mM stock solution in DMSO and cells were dosed at a concentration of 200nM at time 0 and again after 6 hours during the t/baf treatment time course. Bortezomib (bort) (LC Labs #B-1408) was prepared as a 1 mM stock solution in DMSO and cells were dosed at a concentration of 1µM. DMSO was added to control cells at a concentration of 0.02% (v/v).

### Adapting adherent cells to suspension culture

Cells were adapted from ACC to SCC media in a stepwise process of supplementing ACC media with 25%, 50%, 90% and 100% SCC media at each passage. After adaptation, cells were trypsinized and transferred to a 125mL flask for growth in suspension on a shaking platform (Thermo Scientific) at 150 rpm in 95% O_2_, 5% CO_2_. The cells were maintained at a concentration range of 3×10^5^-2×10^6^ cells/ml and kept at a minimum volume of 30mL. Trypsin (Thermo Fisher Scientific) was added (0.008% v/v) to SCC media to reduce cell clumping. Cell growth was monitored using a hemocytometer, where untreated cells grown in suspension cell culture exhibited an average doubling time of 24 hours.

### Pulsed SILAC time course

To initiate each time-course, cells grown in 30mL light-labeled medium were pelleted at 200g for 5 minutes, washed once with a heavy-labeled medium (10mL) and then resuspended in heavy-labeled media bearing either DMSO, torin, t/baf, t/bort, or t/baf/bort. Note that bafilomycin was re-administered in the t/baf time-courses after 6 hours as we had previously observed a time-dependent decline in its activity (**Supplementary Figure 7**). To allow for complete depletion of preexisting pools of ‘light’ amino acids, samples were collected from each time-course and analyzed via mass spectrometry to identify the timepoint corresponding to maximal light protein abundance, and this point was defined as the 0 minute timepoint for each timecourse. This point corresponded to 30 minutes, 1 hour, 2 hours, and 2 hours for DMSO, torin, t/ baf, t/bort and t/baf/bort, respectively. At each timepoint for all timecourses, a 1mL sample was aspirated from the ∼30mL culture, cells were pelleted via centrifugation at 1,500g for 2 minutes, the supernatant was removed, and cells were flash frozen in liquid nitrogen. Additionally, to assess cell viability during the timecourses, we stained cells with trypan blue and counted viable cells using a hemocytometer (**Supplementary Figure 8**), and assessed total protein abundance by calculating RA_L+H_ at each timepoint (**Supplementary Figure 9**).

### Mass spectrometry sample preparation reagents

The following reagents were used for peptide purification: S-trap™ micro columns (Profiti); sodium dodecyl sulfate (SDS, Sigma Aldrich #436143); triethylammonium bicarbonate (TEAB, Sigma Aldrich #18597); 85% HPLC-grade phosphoric acid (Fisher Chemical); HPLC-grade methanol (Fisher Scientific); LCMS-grade acetonitrile and water (VWR); sequencing grade modified trypsin (Promega #V5113); 1,4-dithiothreitol (DTT, Thomas Scientific #C988Q39); iodoacetamide (IAA, CAS: 144-48-9, Fluka # 57670).

### Peptide purification

Samples were processed and digested on the S-trap™ micro columns following the manufacturing protocol with the following alterations. Samples were resuspended and lysed in buffer SL (50mM TEAB, 5%w/v SDS, pH 8.5). A reference standard of HeLa cells grown in medium-labeled culture media, which was lysed in the same buffer, was spiked into each timepoint. Samples were reduced with 10mM DTT at room temperature for 30 minutes and alkylated in the dark with 10mM IAA for 10 minutes at room temperature. Trypsin was resuspended in 50mM TEAB and added to the column at a1:10 trypsin-to-protein ratio by mass and incubated in a 42°C water bath for 2 hours. Digested peptides were dried using a speed vacuum concentrator centrifuge (Savant) and resuspended in MS sample buffer (4% acetonitrile, 0.1% formic acid).

### Mass spectrometry data acquisition

All tryptic peptides were analyzed by liquid chromatography-mass spectrometry (LC-MS) using an Ultimate 3000 UHPLC system (Thermo Scientific) and Q-Exactive HF-X mass spectrometer (Thermo Scientific) or Orbitrap Exploris 480 mass spectrometer (Thermo Scientific). Peptides (∼2μg) were injected onto a PepMap100 C18 precolumn at 300 nL/min in 98% buffer A (water with 0.1% v/v formic acid) and 2% buffer B (acetonitrile with 0.1% v/v formic acid). Peptides were resolved on a PepMap C18 column (75 μm ID x 2cm, particle size 3 μm pore size 100Å; Thermo Fisher Scientific) using a 135-minute linear gradient from 4%-to-30% buffer B. Peptides were ionized by nano spray electrospray ionization and analyzed on the mass spectrometer. For data dependent acquisition runs, precursor (MS^1^) full scans (350-1400 m/z range) were collected at a resolution of 60,000 (AGC 3×10^6^, 50ms maximum injection time). HCD fragmentation of the top 12 abundance precursor ions were performed at 25% normalized collision energy (NCE). Fragment (MS^2^) analysis was performed with a resolution of 15,000 (AGC 1×10^5^, 100ms maximum injection time, 2.2m/z isolation window). For data independent acquisition runs, MS^1^ full scans (350-1400m/z range) were collected at a resolution of 120,000 (AGC 3×10^6^, 50ms maximum injection time). MS^2^ scans were subdivided into 25 DIA isolation windows of varying widths using a resolution of 30,000 (AGC 1×10^6^, 70ms maximum injection time). HCD fragmentation was performed at 25% NCE.

### Mass spectrometry data preprocessing

A protein search was conducted in Spectronaut Pulsar v.15/v.16 (Biognosys AG, Switzerland) on DIA time course samples. DirectDIA workflow, a library-free method, was used for the search with the human protein database (UP05640) as the background proteome. For the t/baf/bort datasets, a hybrid search library was generated using both the DIA and DDA datasets. The default search settings were modified as follows: 1) include 3 label channels in the search (default, [Arg^6^+Lys^6^], [Arg^10^+Lys^8^]); 2) in the Pulsar search, we use the label workflow with *in-silico* generation of missing channels. Protein copy numbers reported following Wisniewski, Hein et al. 2014.

### Data filtering and normalization

Peptides were filtered for quality based on the reference [Arg^6^+Lys^6^] channel using Spectronaut measures of: normalized cscore, signal-to-noise, and peak shape quality score. Following the initial quality filtering, the peptides were manually inspected and curated before proceeding with downstream analysis. Quantitation of each isotope channel was performed at the MS^1^ level by integrated area-under-the-curve for mass isotopomers M_0_, M_+1_, M_+2_, M_+3_ peaks for light labeled precursors, M_0_, M_+1_ peaks for medium labeled precursors and M_+1_, M_+2_, and M_+3_ peaks for heavy labeled precursors. The total abundance was calculated by the addition of light [M_+1_, M_+2_, M_+3_] and heavy [M_+1_, M_+2_, M_+3_] peaks. All light and heavy precursors were normalized to the reference standard following *eq. 1* and *eq. 2*, providing a relative peptide abundance.

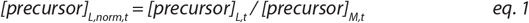

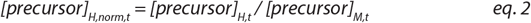

The protein abundance was reported as the median peptide value across mapped peptides. Only proteins with 2 or more uniquely identified peptides were reported.

To analyze kinetic profiles, data from all replicates were pooled and kinetic analyses for protein synthesis, degradation and change in total abundance were performed by calculating the protein abundance relative to t_0_ (RA) (*eq. 3 – 5*).

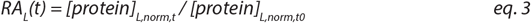

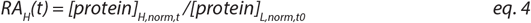

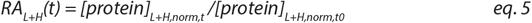

The relative abundance at each time point was normalized by the deviation observed in global total abundance as follows. First, the total abundance was fit to a single exponential (eq. 6) using robust regression to account for outliers, with amplitude and rate parameters *A*, and *k*.

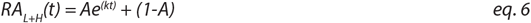

The global total abundance and calculated fits are shown in **Supplementary Figure 9**. Next, the deviation normalization factor is calculated as the ratio of the fit value at time t ([R_AL+H_]_calc,t_) to the median observed value [RAL+H]_obs,t_, following *eq. 7*.

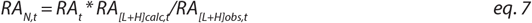

### Kinetic modeling of protein synthesis

We measured protein synthesis by modeling the change in heavy isotope incorporation over time. To account for potential variations in translational capacity resulting from cell growth and division in untreated cells, we first normalized the measured relative abundance at each time point using the median total abundance of the large ribosomal subunit, as determined by quantitative mass spectrometry (qMS).

In modeling protein dynamics, we assumed that protein synthesis was a zeroth order process, and protein degradation was a first order process. We represented the rate at which heavy isotopes are incorporated following *eq. 8*.

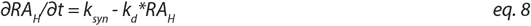

Integration of *eq. 8* results in the integrated rate equation (*eq. 9*), which encapsulates the synthesis (k_s_) and degradation (k_d_) of nascent proteins.

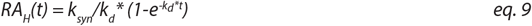

### Kinetic modeling of protein degradation

Kinetic modeling followed the workflow depicted in **Supplementary Figure 1**. Protein degradation profiles were fit using 1-phase (2-parameter) and 2-phase (4-parameter) models of protein degradation, as outlined in equations 10-11.

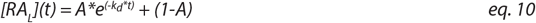

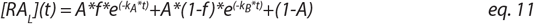

Following eq. 11, the amplitudes for phase A and B are expressed as:

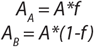

The half-life from *eq. 10* and *eq. 11* can be expressed as:

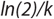

### Kinetic model selection

All datasets were analyzed using Prism version 10.1.0, with 1-phase or 2-phase degradation kinetic models selected based on the Akaike Information Criterion (ΔAICc, *eq. 12*) (Akaike 1973), with a ΔAICc > 2 resulting in the 2-phase degradation model selection (*eq. 11*), and those ΔAICc < -2 resulting in the 1-phase model selection (*eq*.*10*). These cutoffs correspond to ∼73% model likelihood.

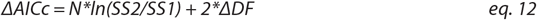

where SS2 and SS1 refer to the sum squared residual of the 2-phase and 1-phase fits, respectively; N references the number of datapoints; and *ΔDF* refers to the change in the number of free parameters between the two models (2, in this instance).

We evaluated the fit quality by the multi-parameter root mean square error, Sy.x (*eq. 13*).

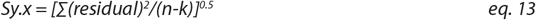

where n is the number of data points and k is the number of parameters used in the curve fitting. Peptides with Sy.x>0.15 and those with fewer than 4 data points were eliminated from further analysis.

### Sucrose density gradient sedimentation assay

HeLa cells were treated with DMSO or 100mM torin for 18 hours, rinsed with cold PBS and then lysed in buffer HLB (20mM Tris, 150mM NaCl, 5mM MgCl_2_, 1% Triton X-100, 1mM DTT, and 0.1mg/ml cycloheximide, pH 7.3). The cell lysates were centrifuged at 21,000g for 5 minutes to pellet the cellular debris. The RNA concentration in the supernatant was quantified by measuring absorbance at 260nm, and the RNA concentration as normalized across samples via dilution in buffer HLB. The samples were layered onto a prepared sucrose gradient, ranging from 10% to 50% (w/v) sucrose in a buffer PB (20mM HEPES, 5mM MgCl_2_, 100mM KCl, 1mM DTT, and 0.1mg/ml cycloheximide, pH 7.3). The gradients were subjected to ultracentrifugation using a SW41 TI rotor (Beckman) at 21,000g for 2.5hours at 4°C. Post-centrifugation, the gradients were analyzed with a piston gradient fractionator (BioComp) by monitoring absorbance of the eluate at 260 nm.

## ACKNOWLEDGMENTS

We thank Bertina Telusma, Jen Kosmatka, Laurel Kinman, April Lee, Alicia Darnell, Joel Sexton and Jennifer Chu helpful discussions, and Martin Taylor, Skylar Kim, Malene Hansen, Terje Johansen, Rich Schiavoni, and Jon Zmuda and technical guidance and reagents. The Davis lab is supported by NIH grants R01-GM144542, R00-AG050749, and NSF-CAREER grant 2046778.

## CONFLICTS OF INTEREST

The authors declare no conflicts of interest.

## CONTRIBUTIONS

DSC and JHD designed the project. DSC and SMW prepared the samples and collected the data. DSC and JHD performed all the analyses, prepared figures, and wrote the manuscript. All authors were involved in preparing the final draft of the manuscript.

## DATA AND SOFTWARE AVAILABILITY

The correspondence for material and data in the manuscript should be addressed to The raw mass spectrometry data and summary tables will be publically deposited upon publication. An interactive data browser is available at jhdavislab.org/datasets.

## SUPPLEMENTARY INFORMATION

**Supplementary Figure 1.**
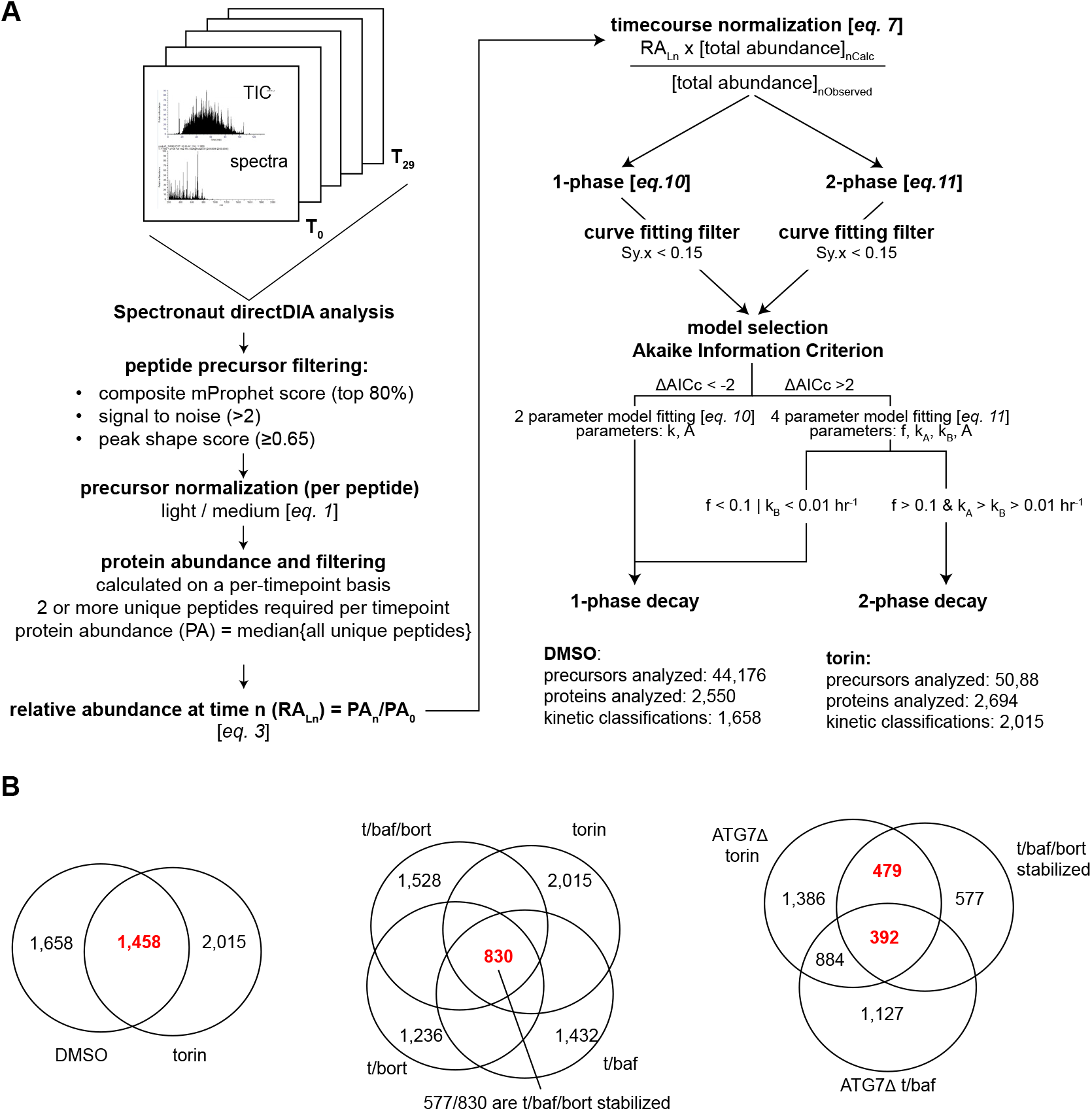
Pulsed SILAC quantitative mass spectrometry data processing workflow. **(A)** Workflow depicting data filtering, normalization, and kinetic modeling approaches employed (see Methods for a more detailed description). **(B)** Quantitation of number of proteins passing all analysis and modeling filters in each condition. Key protein intersections described in this work highlighted in red.

**Supplementary Figure 2.**
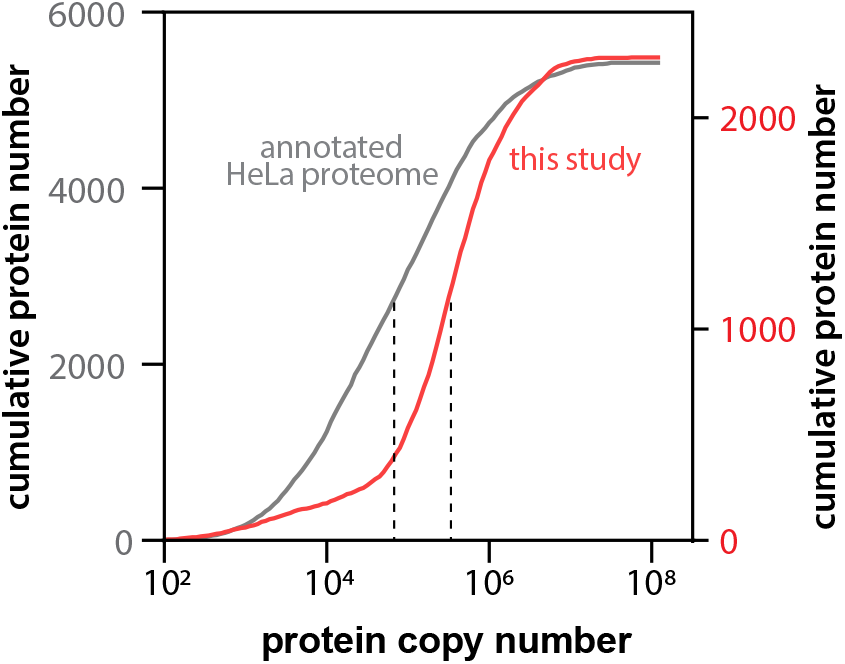
Distribution of protein copy number of detected proteins. Cumulative distribution plot of the protein copy number within the HeLa proteome as estimated previously (Wisniewski et al., 2014) on the left axis (grey) and proteins quantified in this study on the right axis (red). The median copy number for each distribution is marked with a dashed line.

**Supplementary Figure 3.**
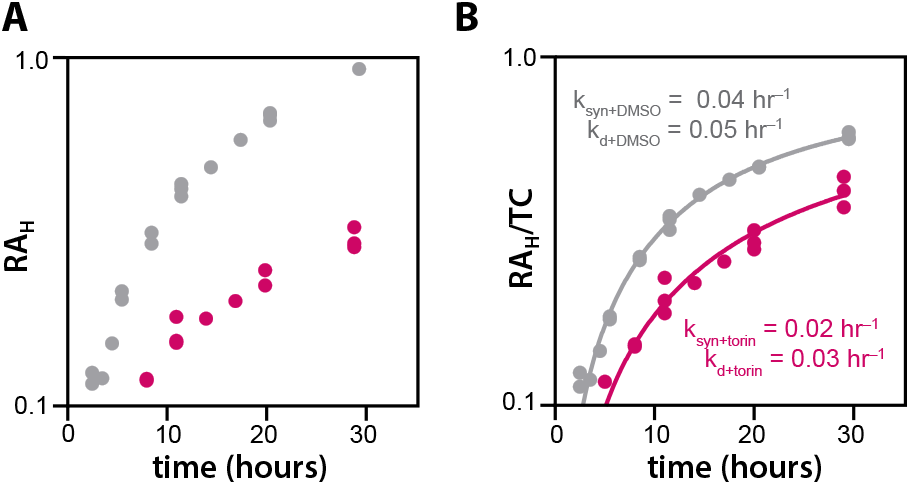
Torin treatment downregulates protein synthesis globally. **(A)** The median relative post-pulse (*i*.*e*. heavy isotope labeled) protein abundance across all proteins plotted on a semi-log axis for torin-treated (pink) and control (grey) conditions. **(B)** Data from A normalized to the median total abundance of the large ribosomal subunit at each time point. This normalization was performed to account for growth-dependent changes in total translational capacity. In each condition, data were fit to *eq. 9* (see Methods), and fit kinetic parameters for synthesis (k_syn_) and degradation (k_d_) of nascent proteins are noted.

**Supplementary Figure 4.**
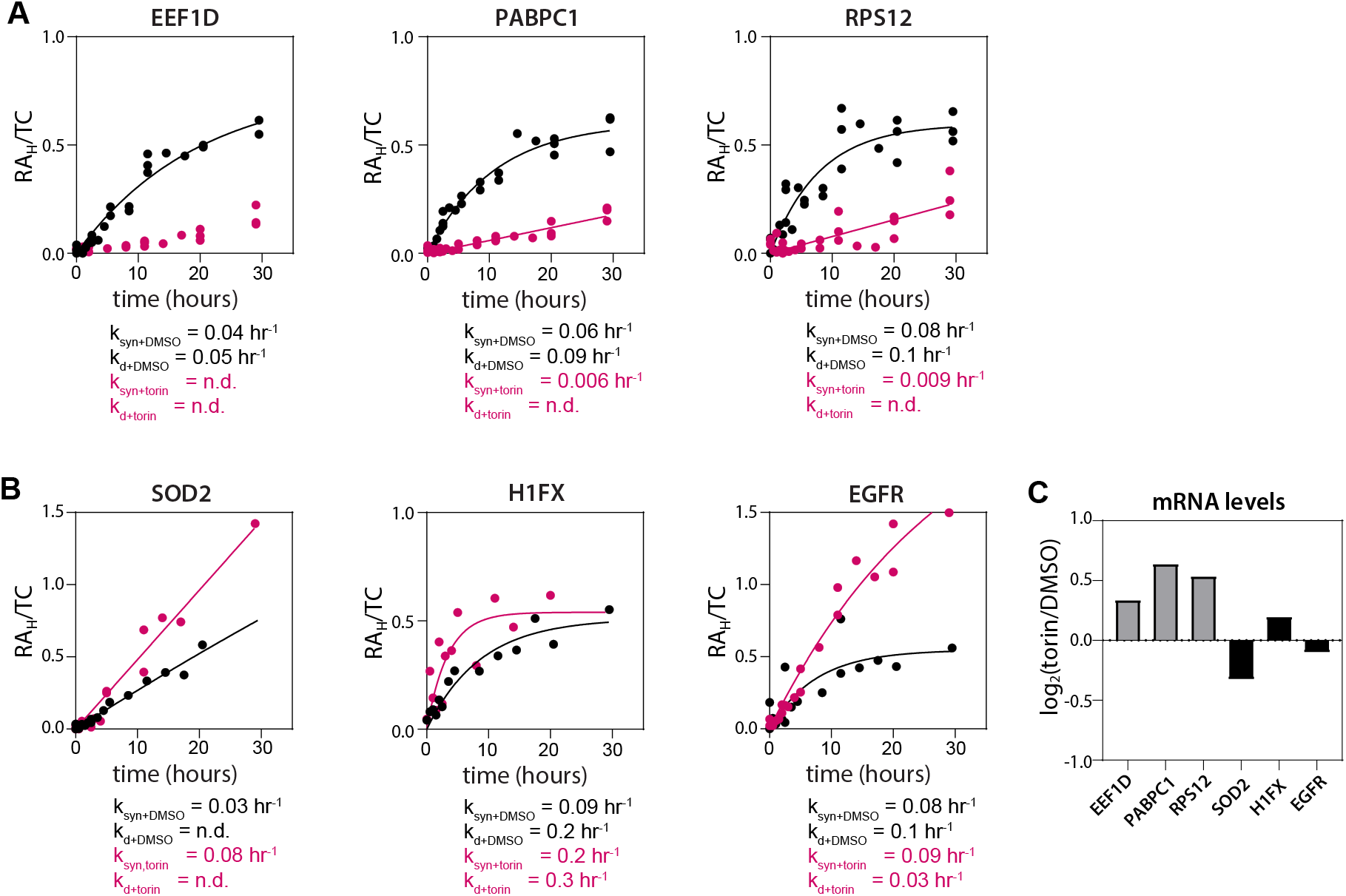
Torin selectively inhibits the synthesis of 5’TOP motif bearing mRNAs. **(A)** Exemplar kinetic traces of protein synthesis corrected for translational capacity (RA_H_/TC) for proteins bearing 5’TOP motifs in their transcripts under torin treatment (pink) and control conditions (black). The data were fit to *eq. 9*, with the fit protein synthesis (k_syn_) and degradation (k_d_) rates listed. **(B)** Exemplar traces and fits for proteins whose synthesis was not repressed by torin treatment, analyzed as in (A). **(C)** mRNA expression levels for the proteins noted in panels (A-B), extracted from analysis of HEK293T cells following 24-hour torin treatment (Park, Reyna-Neyra *et al*. 2017).

**Supplementary Figure 5.**
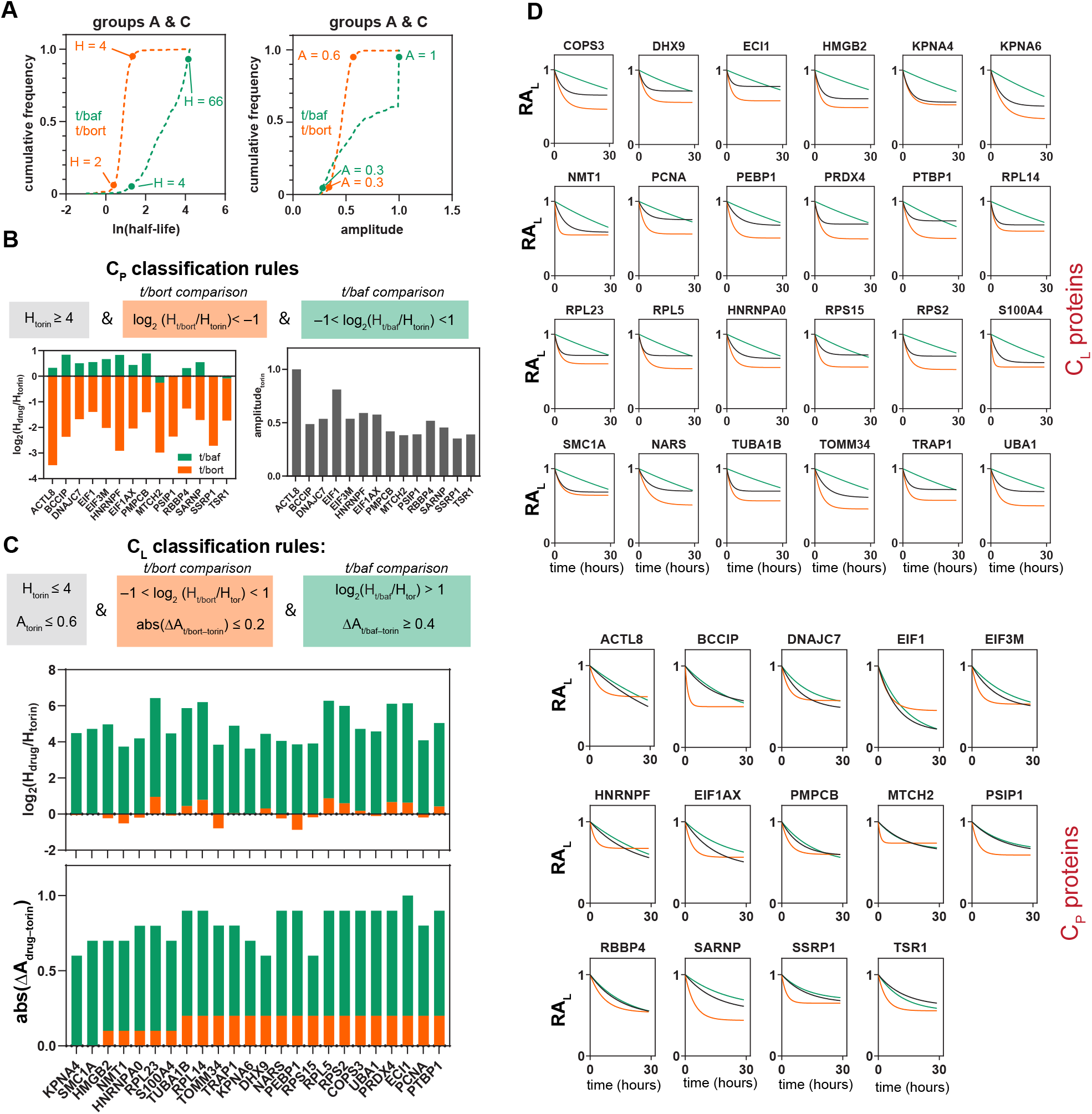
Classification of compensatory proteasomal and lysosomal substrates. **(A)** Cumulative frequency plots of kinetic parameters for group A and C proteins under t/baf (green) and t/bort (orange) conditions. The 5^th^ and 95^th^ percentile values are noted on the plots. **(B)** Classification rules for compensatory proteasomal (C_P_) proteins (top), and bar chart plotting the measured kinetic parameters for C_P_ assigned proteins (bottom). Changes in kinetic parameters relative to those measured in torin-treated cells are colored green (t/baf) and orange (t/bort). **(C)** Classification rules for compensatory proteasomal (C_L_) proteins (top), and bar chart plotting the measured kinetic parameters for C_L_ assigned proteins (bottom). Bar charts are colored as in (C). **(D)** The kinetic fits of proteins assigned to the C_L_ (top) and C_P_ (bottom) groups in cells treated with torin (black), t/baf (green), and t/bort (orange).

**Supplementary Figure 6.**
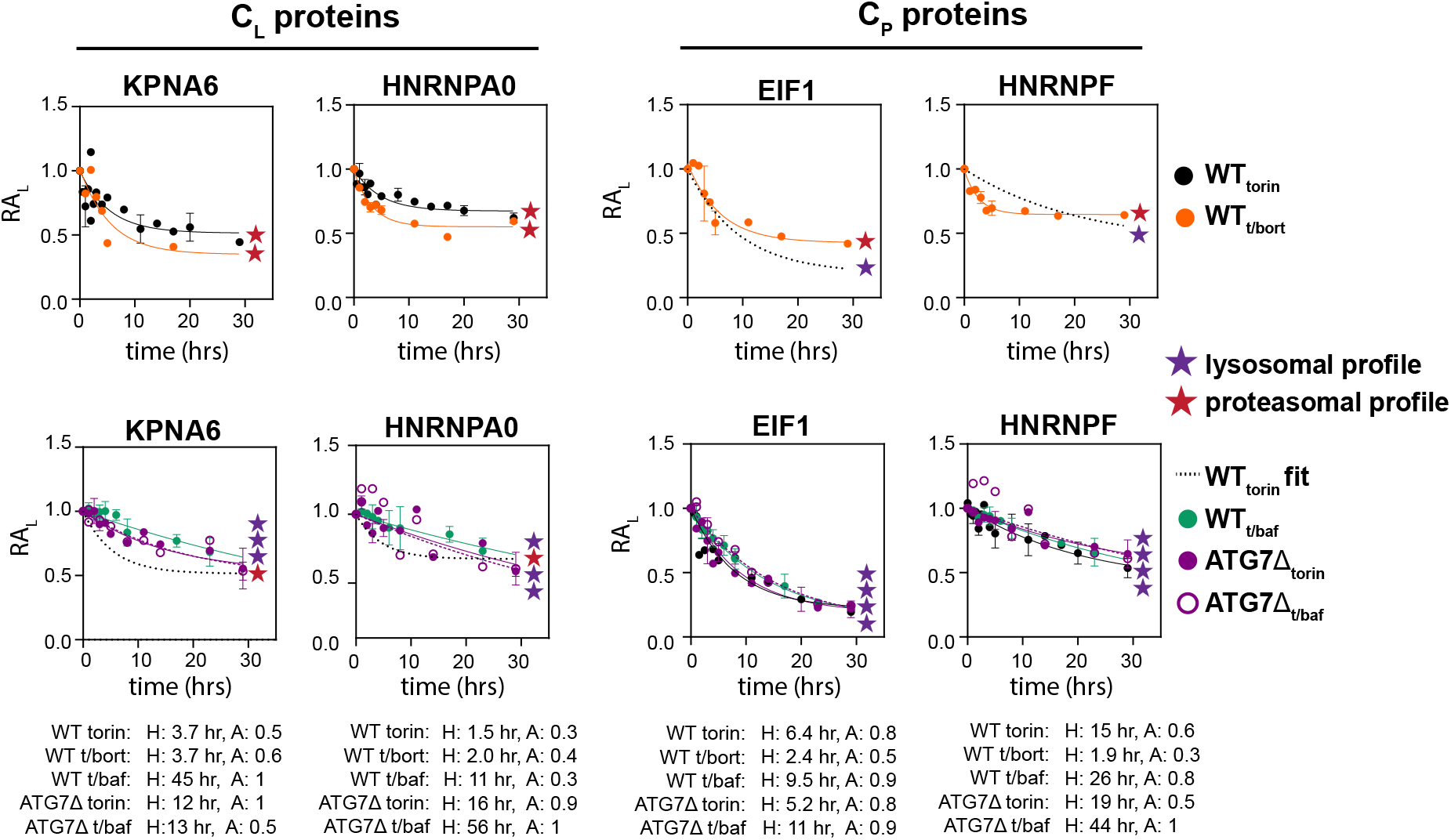
Group C_L_ proteins exhibit proteasomal degradation kinetics in ATG7Δ cells. Exemplar kinetic traces of proteins from group C_L_ (left) and C_P_ (right) in wild-type or ATG7Δ cells treated with torin, t/baf, or t/bort. Profiles characteristic of lysosomal and proteasomal degradation are noted with purple and red stars, respectively. Each time course was fit to *eq. 10*, with kinetic parameters listed.

**Supplementary Figure 7.**
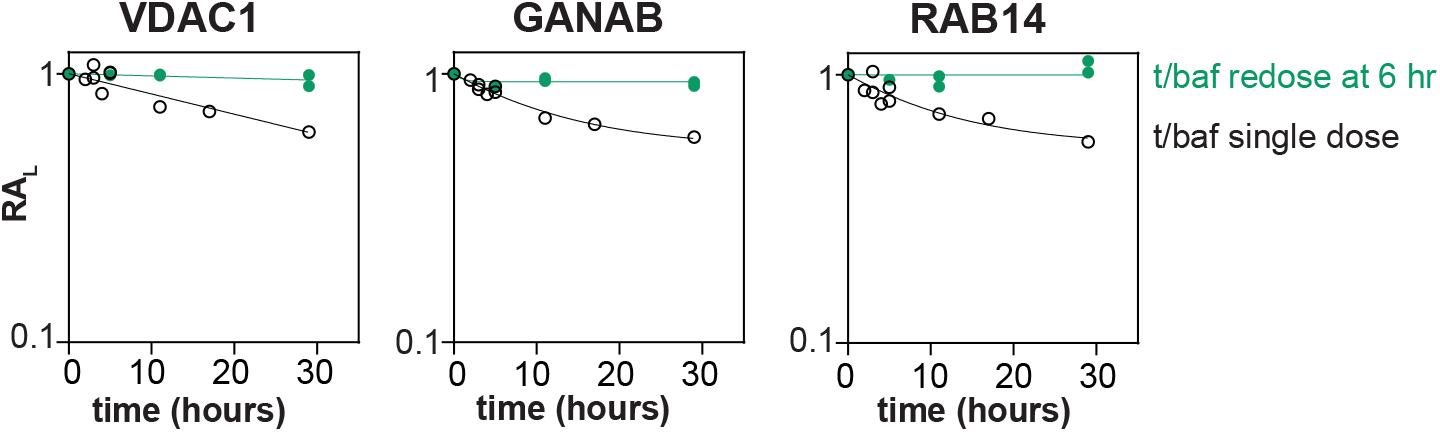
Re-dosing with bafilomycin A stabilizes lysosomal substrates. Kinetic traces of exemplar lysosomal substrates either treated with a single dose of bafilomycin A (black), or re-dosed after 6 hours (green).

**Supplementary Figure 8.**
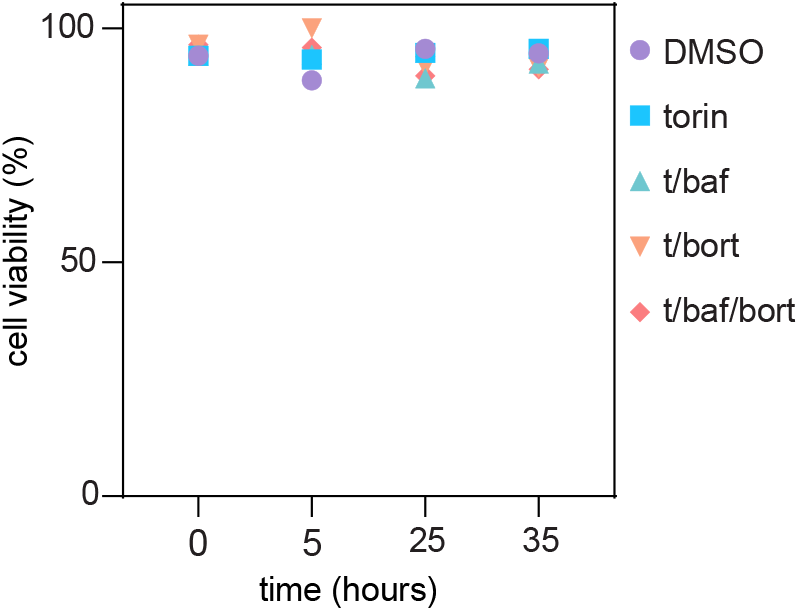
Suspension HeLa cells maintain viability throughout time-course. Cell viability was determined for cells treated with DMSO, torin (100 nM), t/baf (200 nM baf), t/bort (1 µM bort), or t/baf/bort co-treatments using the same concentrations. At each timepoint, cells were aspirated, pelleted at 400g and resuspend with PBS. Cells were mixed with trypan blue at a 1:1 ratio and staining was quantified using a hemocytometer. Percentage of unstained cells plotted as cell viability.

**Supplementary Figure 9.**
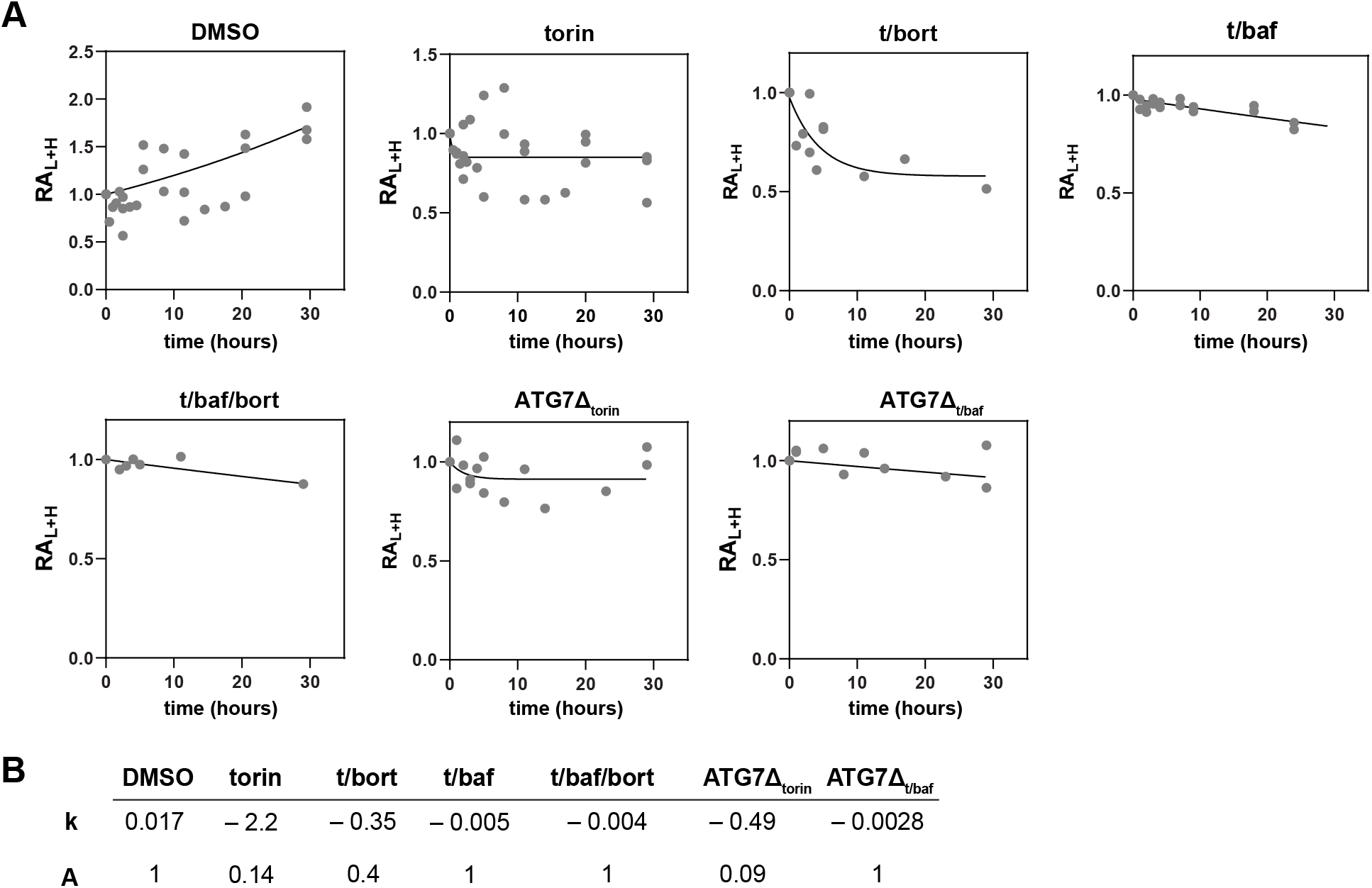
Kinetic fits used for data normalization. **(A)** The median total relative abundance (RA_L+H_) at each time point across the proteome is plotted. For each condition, these data were fit to *eq. 6*. **(B)** Fit parameters from (A) used to calculate [RA_L+H_]_calc,t_ for each experimental condition (see Methods).

## Notes

### Competing Interest Statement

The authors have declared no competing interest.

